# Modeling tissue-specific breakpoint proximity of structural variations from 2,382 whole-genomes to identify cancer drivers

**DOI:** 10.1101/2021.09.27.461957

**Authors:** Alexander Martinez-Fundichely, Austin Dixon, Ekta Khurana

## Abstract

Structural variations (SVs) in cancer cells often impact large genomic regions with functional consequences. However, little is known about the genomic features related to the breakpoint distribution of SVs in different cancers, a prerequisite to distinguish loci under positive selection from those with neutral evolution. We developed a method that uses a generalized additive model to investigate the breakpoint proximity curves from 2,382 whole-genomes of 32 cancer types. We find that a multivariate model, which includes linear and nonlinear partial contributions of various tissue-specific features and their interaction terms, can explain up to 57% of the observed deviance of breakpoint proximity. In particular, three-dimensional genomic features such as topologically associating domains (TADs), TAD-boundaries and their interaction with other features show significant contributions. The model is validated by identification of known cancer genes and revealed putative drivers in novel cancers that have previous evidence of therapeutic relevance in other cancers.

## Introduction

Whole-genome sequencing of cancer genomes has revealed that they contain a wide variety of DNA structural variations (SVs) that include deletions, duplications, translocations and other complex events [1]. The SVs in cancer cells arise from different mechanisms and vary in size from kilobases to whole chromosomal rearrangements [1-4]. Consequently, SVs usually span several genes and their associated regulatory elements. While it is well known that genomic rearrangements and copy number variations (CNVs) can lead to dysregulation of tumor-suppressors or oncogenes and act as drivers of cancer progression [1, 5-8], identification of SVs under positive selection in cancer remains a challenging task. This is because SVs are heterogeneously distributed across the genome leading to many genomic regions recurrently altered in multiple samples due to neutral background processes [4]. To identify the events under positive selection, the null background distribution of SV breakpoints must be characterized accounting for the genomic covariates [5, 9]. Additionally, the identification of the specific functional element that is the target of positive selection (i.e., the coding sequence of a gene, its cis-regulatory regions, or non-coding RNAs) constitutes another challenge due to the large genomic span of SVs.

While numerous computational methods have been developed to model the background distribution of single-nucleotide variants (SNVs) and identify drivers in a tissue-specific manner, similar methods for SVs are lacking [5, 10-13]. The Pan-Cancer Analysis of Whole Genomes (PCAWG) SV Working Group used a Gamma-Poisson fit to model the breakpoint density from 2,658 whole-genomes using eight covariates to identify the significant driver genes in several cancers [5]. However, this analysis was performed at the pan-cancer level without accounting for tissue-specific covariates. Since there is ample evidence of different SV distributions and putative mechanisms across cancer types [1, 2, 14], it is critical to model SV breakpoint distribution in a tissue-specific manner to obtain the corresponding accurate null models. Importantly, the three-dimensional (3D) high-order genomic structure, such as the topologically associating domains (TADs), has not previously been considered a covariate of breakpoint distribution. Recent studies have analyzed the relationship between 3D topology and SVs in cancer [8, 15-17]. It was reported that while SVs can lead to changes in chromatin folding, only 14% of TAD-boundary deletions are associated with significant gene expression changes [16]. This finding indicates that many of these SV events may be related to neutral evolution in tumor cells and must be accounted for in the null model to identify true drivers. Furthermore, previous studies did not account for the non-linear contribution of covariates or their interactions in the modeling of SV breakpoint distribution.

Here, we describe a new method that models the breakpoint proximity of SVs in 2,382 cancer whole-genomes in a tissue-specific manner. We used nine different genomic covariates, including the recurrence of TADs and TAD-boundaries across cell lines and tissues as well as their functional classification based upon chromatin states. We implemented a generalized additive model (GAM) to describe the genome-wide SV breakpoint proximity. The use of a GAM allows us to include both the linear and non-linear contributions of variables as well as their interplay. Modeling breakpoint proximity has the advantage of allowing us to analyze the clustering of breakpoints over dynamic genomic lengths to capture different breakpoint trends, unlike breakpoint-density-based approaches that rely on fixed genomic bin sizes. Finally, we use this approach to identify loci that exhibit signals of positive selection and the functional elements that are the likely targets of selection. The method is implemented in Cancer Structural Variation Drivers (CSVDriver), a user-friendly tool that can be used by researchers to model the SVs from whole-genome sequencing data and identify putative cancer drivers.

## Results

We analyzed a set of 324,838 high confidence somatic SVs derived from whole-genome sequencing of 2,382 patients of 32 distinct cancer types from 15 different organ systems (Supplementary Table 1). The cancer types include those with a high SV burden from the PCAWG project [1, 18] and metastatic prostate cancer samples [19, 20]. Based on the rationale that tissue-specific covariates can influence the rearrangement landscape, the cancer types from different organ systems were modeled separately. Furthermore, the prostatic primary and metastatic cohorts were analyzed separately since they can have distinct drivers.

### Breakpoint proximity curve to model genome-wide SV distribution

To describe the genome-wide distribution of SV breakpoints in a given cohort, we included all unique breakpoint coordinates for each sample. Then we computed the breakpoint proximity curve (*BPpc*) based upon the breakpoint neighbor reachability (*BPnr*_*i*_) for each individual breakpoint (*BP*_*i*_) in the cohort. This metric captures the genomic regions with high or low proximity between breakpoints (Methods). The *BPpc* is defined as the smooth curve resulting from the nonparametric local polynomial regression (locally estimated scatterplot smoothing, LOESS) fitted to the dataset of *BPnr*_*i*_, after reverse scale normalization *-log10(BPnr*_*i*_*+1)* (Fig. 1a). *BPpc* allows the use of a peak-calling approach to directly identify the regions with higher breakpoint clustering relative to the surrounding area (i.e., peak summits), thereby overcoming the inherent issues associated with pre-defining a genomic bin size for computing breakpoint density along the genome [5, 21]. This is important because functionally relevant breakpoint clustering events may occur over a wide range of genomic lengths. Thus, *BPpc* models the underlying distribution of breakpoints in a more robust and unsupervised manner compared to the computation of breakpoint densities.

**Figure 1:**
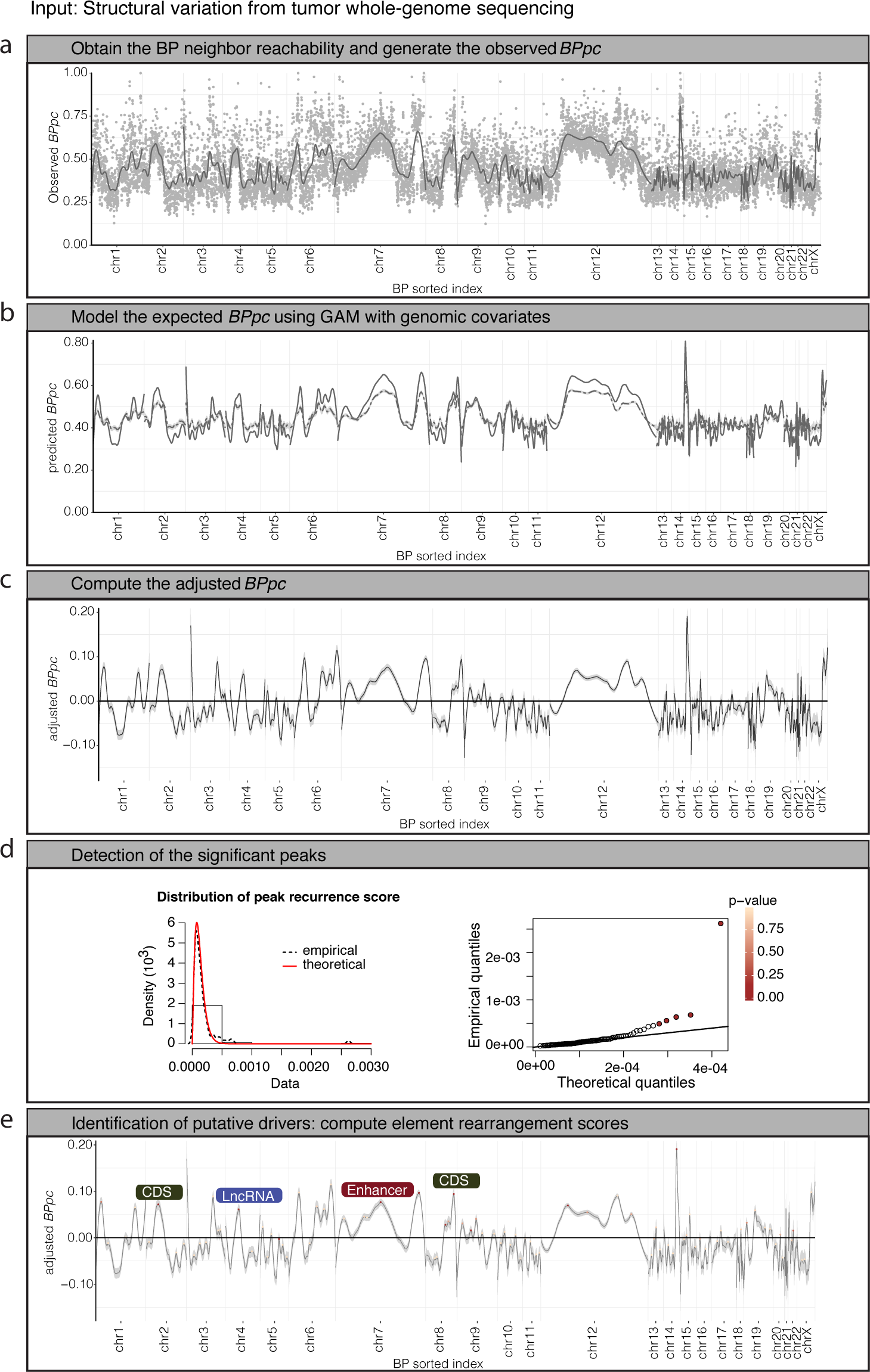
CSVDriver workflow. The input data is the standard cancer somatic SV calls that include a pair of breakpoint coordinates (BP1, BP2) for each SV_id per sample. **(a)** Step 1 computes the breakpoint neighbor reachability (BPnri grey dots) for each breakpoint (*BP*_*i*_), where ‘*i*’ is the sorted index. Then, for each chromosome, the *BPpc* is generated as the smooth curve (grey line) resulting from the nonparametric local polynomial regression fitted to the dataset of *BPnr*_*i*_, after reverse scale normalization *-log10(BPnr*_*i*_*+1)*. **(b)** In step 2, based on the observed *BPpc* distribution (solid line), the method assesses the expected 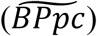 background distribution (dashed line) by using a generalized additive model (GAM) that includes multiple tissue-specific breakpoint covariates. **(c)** In step 3, it computes adjusted *BPpc* (observed – expected). **(d)** In step 4, the method calls peaks across the adjusted *BPpc* and identifies those that potentially correspond to positively selected loci. It computes the peak recurrence score (PRs) and based on the empirical density of PRs (dashed line distribution) it identifies the peaks with PRs significantly higher (QQ-plot FDR < 0.2) than the fitted theoretical density (red line distribution). **(e)** In the last step 5, the driver candidates are identified as the genomic elements (CDS (coding sequence), enhancer, CTCF-Insulator, and lncRNA) with the highest rearrangement scores within the peak region.

### Expected *BPpc* using genomic covariates in a GAM

The core of the genomic breakpoint distribution arises under neutral selection due to background processes likely corresponding to the tissue-specific functional and structural genome heterogeneity. We modeled the expected background *BPpc* per tissue type using a GAM with multiple variables (Supplementary Table 2), including the tissue-specific chromatin state annotations from the Roadmap Epigenomics project [22] [23], recurrence of TADs (TAD.recurr), and TAD-boundaries (TAD-B.recurr) from the 3D Genome Browser [24], replication timing (RT) from ENCODE Repli-seq data [25] and fragile sites (FS) from the HumCFS database [26]. Other variables include the genome complexity from UCSC genome browser repeat elements data [27], gene density and GC content (GC). Thus, the expected breakpoint proximity curve is defined as 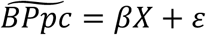, a variation of the generalized linear model in which X is a matrix of covariates (breakpoint predictors) that can contribute linearly as well as non-linearly to the response variable *BPpc* (Methods) (Fig. 1b).

The multivariate model explained a larger proportion of the null deviance compared to the single predictor models, even without including the interplay of co-variates (Fig. 2a). The median explained deviance is 18% and ranges from 6% for lung cancer to 50% for lymph-nodes (Fig. 2a). We expected that modeling the interaction effects of genomic features may contribute substantially to the model due to the underlying complexity of SV breakpoints in cancer [21]. Since all possible combinations of features make the model computationally infeasible and intractable for a meaningful interpretation, only the interaction terms of features that contribute at least 10% in at least one cohort in single predictor models were included (Fig 2a). The resulting model accounts for the interaction of various features, including RT, GC, TAD.recurr, TAD-B.recurr and gene density, with lamina-associated domains (LADs) that correspond to the higher-order genome disposition within the nucleus [28]. The model also accounts for the pairwise interplay between gene density and RT, gene density and TAD.recurr as well as RT and TAD.recurr, in each case the interactions are modeled separately for distinct TAD segment classes. The TAD segments are defined based on differential conservation across cell types and annotated into three classes (quiescent, low-active or active) using enrichment of tissue-specific chromatin annotations [16] (Methods). The multivariate GAM that accounts for the interaction of co-variates showed further substantial improvement over the model without the interaction terms. The median explained deviance is 29%, ranging from 10% in lung cancer to 57% in lymph-nodes (Fig. 2a). The model performs well in all cohorts (Fig. 2c) with fairly narrow and symmetric residual distributions (Supplementary Fig. 1). The explained deviance for each cohort is not biased by the sample size as shown by its lack of correlation with the average number of SVs per donor (Supplementary Fig. 2). Notably, we find a substantial decrease in the correlation of observed vs. predicted *BPpc* when using the covariates from unmatched tissues (Fig. 2b). This clearly highlights the importance of using tissue-specific co-variates.

**Figure 2:**
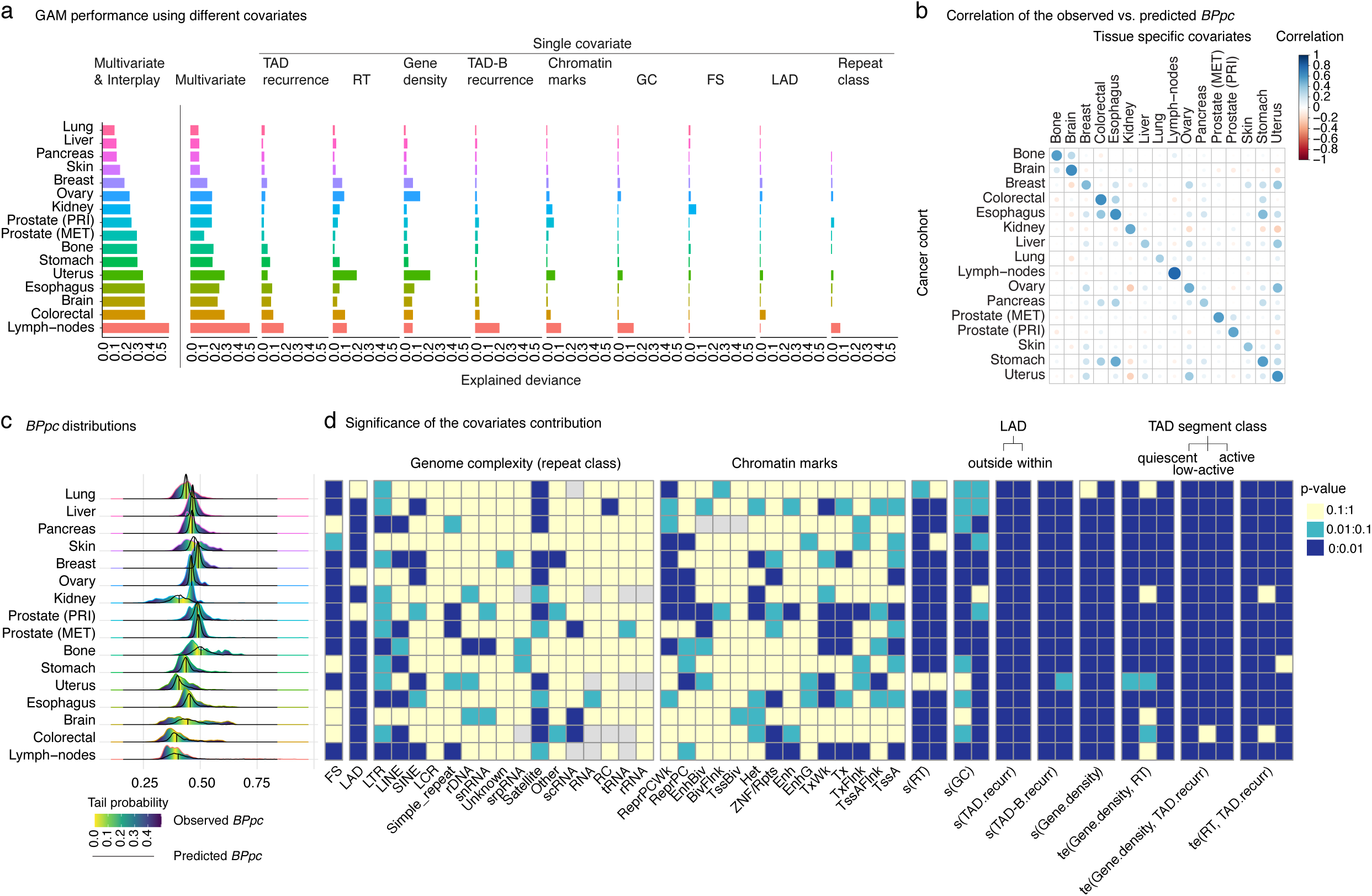
Performance of the GAM for the expected background BPpc for each cancer type, **(a)** Explained deviance of the GAM for the multivariate model with and without interplay and single covariate models, **(b)** Correlation plot of the performance of the model for each cohort using different tissue-specific covariates, **(c)** Observed BPpc distributions and heatmap of the tail probabilities. Black curve is the predicted BPpc distribution from the GAM model. **(d)** Heatmap of the significance of the partial contribution of each covariate to the explained deviance of the model with respect to the mean of the distribution (null intercept). FS (fragile sites), LAD (lamina associated domain), DNA sequence repeat classes that include LTR (long terminal repeat elements), LINE (long interspersed nuclear elements), SINE (short interspersed nuclear elements) including ALUs, LCR (low complexity repeats), simple repeats i.e. micro-satellites, DNA repeat elements (rDNA), RNA repeats (including tRNA, rRNA, snRNA, scRNA, srpRNA), satellite repeats, other repeats including class RC (Rolling Circle) and Unknown complex sequence.chromHMM tissue-specific chromatin marks that include TssA (Active TSS); TssAFlnk (Flanking Active TSS); TxFlnk (transcription at gene 5’ and 3’); Tx (Strong transcription); TxWk (Weak transcription); EnhG (Genic enhancers); Enh (Enhancers); ZNF/Rpts (ZNF genes & repeats); Het (Heterochromatin); TssBiv (Bivalent/Poised TSS); BivFlnk (Flanking Bivalent TSS/Enh); EnhBiv (Bivalent Enhancer); ReprPC (Repressed PolyComb); ReprPCWk (Weak Repressed PolyComb). The ‘s’ is a thin plate regression spline smooth function that describes the non-linearity in the contribution of the replication timing (RT), GC content (GC), gene density (gene.density), and the recurrence of the topologically associated domain (TAD.recurr) as well as the recurrence of TAD boundary regions (TAD- B.recurr). The partial contribution of each ‘s’ function accounts for the interaction with the corresponding status of LAD. The ‘te’ is a tensor product interaction that describes the interplay between gene.density vs. TAD.recurr, gene.density vs. RT, and TAD.recurr vs. RT. The partial contribution of each ‘te’ interplay accounts for the interaction with the corresponding class of TAD-segment that includes quiescent, low-active, and active regions.

### Contribution of genomic covariates to the background *BPpc* distribution, nonlinearity and covariates interplay

We find that the contribution of the chromatin state annotations and repeat elements is limited while the 3D genome features of TAD-recurrence and TAD-B recurrence contribute significantly to the model for most cancer types (Fig. 2d). This result is consistent for both the genomic loci close to the membrane of the nucleus, i.e. -within LAD-, as well as the inner-nucleus genomic regions, i.e. -outside LAD-(Fig. 2d). This suggests that the 3D chromosomal conformation plays an important role in the expectation of the genome-wide distribution of SVs. The results also show that fragile sites, replication timing, GC content and gene density contribute significantly to many cancer types, though the behavior can vary for within vs. outside of LADs for some cohorts (Fig. 2d). The two-dimensional (2D) contour plots allow us to visualize the partial contributions of features to *BPpc*. Most cohorts exhibit peaks and valleys on the smooth function for the partial effects of most covariates demonstrating the importance of non-linear modeling (Figs. 3a, 3b, Supplementary Fig. 3). For instance, the partial contribution of TAD-recurrence shows non-linear behavior in brain, colorectal, kidney and prostate cancers although it is linear in breast cancer regardless of the LAD status (Fig. 3a). The behavior of some features may also vary within vs. outside LADs. For example, the contribution of GC content in breast cancer is linear inside LADs but non-linear outside LADs (Fig. 3b).

**Figure 3:**
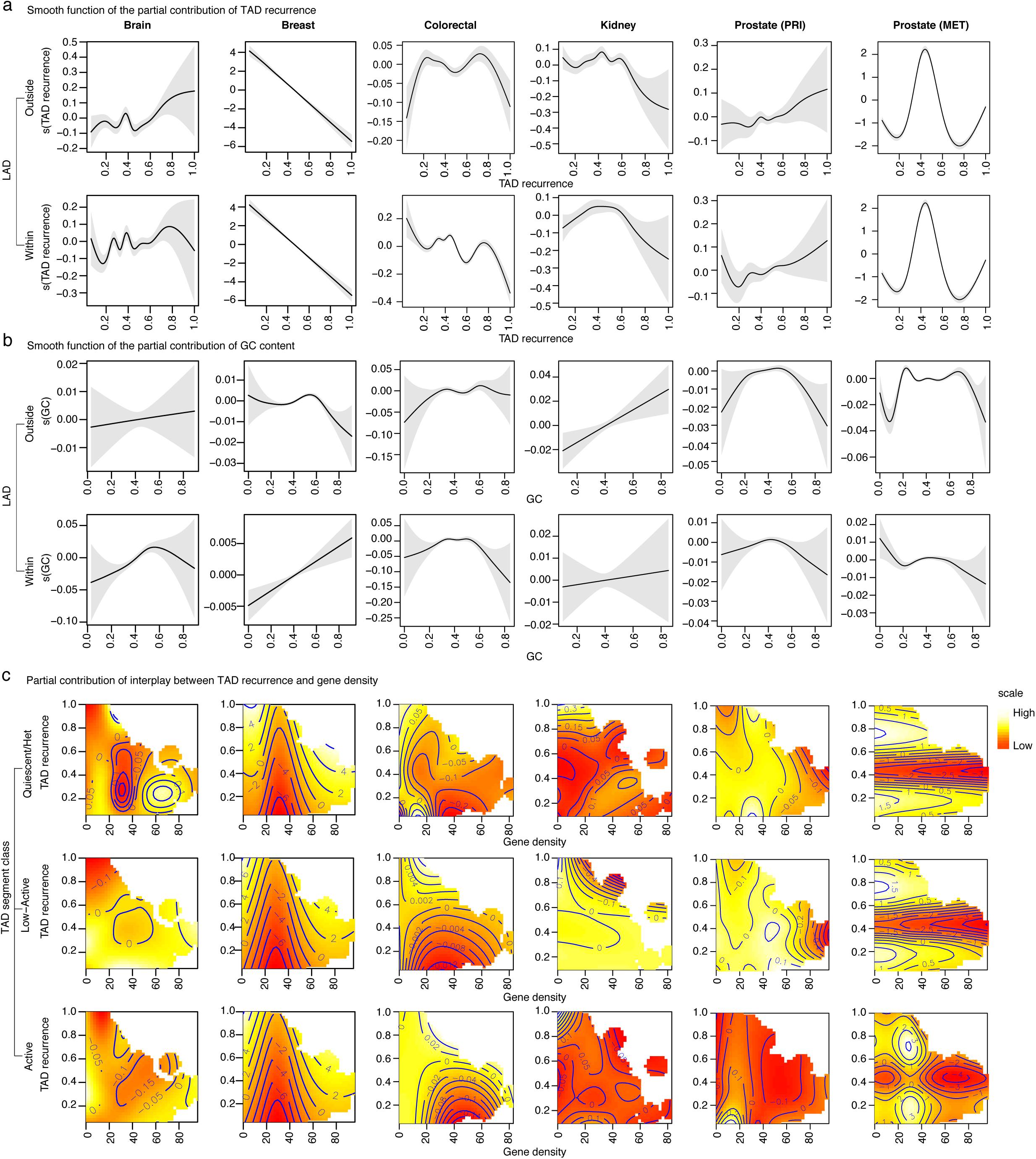
Graphics to analyze the results of the partial contribution of covariates to the GAM shown for representative cancer types. **(a)** the smooth function of partial contribution of TAD recurrence. **(b)** the smooth function of the partial contribution of GC content. These graphics represent the linear or non-linear behavior of the partial contribution of each covariate analyzed. The x-axis is the value of the covariate and the y-axis is the corresponding partial effect of the covariate. The confident interval is represented by the grey area. **(c)** 2D graphics for the partial contribution of the interplay between gene density and TAD recurrence for the interaction with the three functional classes of TAD segments. The scale from light yellow to red represents partial contribution for higher to lower values of the distribution. The full set of graphics for all cohorts is shown in Supplementary Fig. 3

We find a statistically significant contribution for the interaction terms of covariates across all the three functional classes of TAD segments for most cohorts (Fig. 2d). However, their effect sizes show distinct behavior for each cancer type and often across TAD segment classes as evident from the 2D contour plots (Fig. 3c, Supplementary Fig. 3). Interestingly, the distinct behavior is also apparent for primary vs. metastatic prostate cancers, likely pointing to different processes contributing to early vs. late genome-wide SV distribution [29, 30].

In general, there is wide variability in the partial contributions of different features across cancer types demonstrating the importance of tissue-sepcific modeling.

### Putative driver candidates identified at significantly recurrent peaks in *BPpc*

We obtained the adjusted 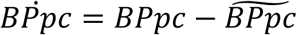 by correcting the observed curve with the expected model (Fig. 1C). Values around zero in the adjusted curve correspond to the observed *BPpc* close to the expected one from the GAM. Peaks in positive values of 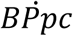 correspond to regions where the breakpoints are closer than expected while the valleys in negative values are loci where the breakpoints are sparser than expected (Fig. 4a and Supplementary Fig. 4). As expected, there is a wide variability of the peaks and valleys representing the differential landscape of rearrangements for each cohort (Fig. 4a and Supplementary Fig. 4)

**Figure 4:**
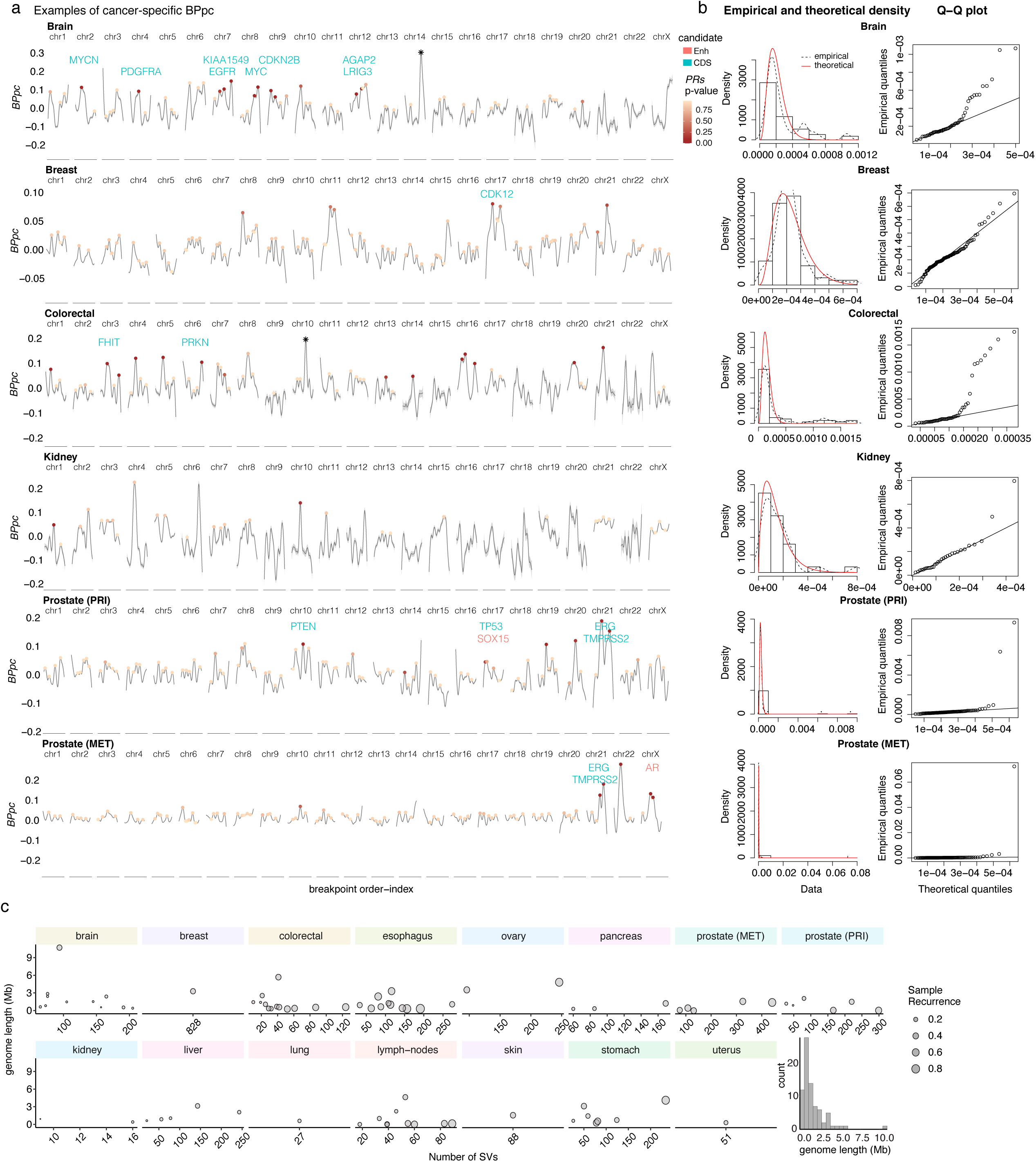
**(a)** Examples of cancer-specific *BPpc* and the peaks of significantly recurrent rearrangements for representative cancer types. Each peak shows a dot colored in the scale of significance for the corresponding peak recurrence score (PRs). The peaks that come mostly from a unique sample (likely a chromothripsis event) are marked with an asterisk. The known driver candidates detected within the significantly rearranged peaks are marked (green for coding sequence and red for enhancers). **(b)** Empirical and theoretical density of PRs for each cohort and the corresponding QQ-plots which show the p-values follow a uniform distribution. **(c)** Scatter plots of the genome length of the peak region vs. number of SVs for each peak, the size of the dots shows the recurrence in the cohort. Histogram of the genome length distribution of the significant peaks.

To identify the peaks that potentially correspond to positively selected loci across the *BPpc* rearrangement landscape, we computed the peak recurrence score (*PRs*) (Methods).

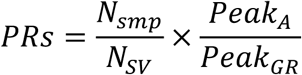

where *Nsmp* is the number of unique samples in the peak, *Nsv* is the number of unique SVs, *Peak*_*A*_ is the area under the peak and *Peak*_*GR*_ is the genome range of the peak. Thus, *PRs* is highest for peaks where many samples (high *Nsmp*) contribute same SVs (low *Nsv*) to create tight clusters of breakpoints (high *Peak*_*A*_) over narrow regions (low *Peak*_*GR*_). Next, for each cohort, we identified peaks with *PRs* significantly higher than the fitted theoretical density using a gamma distribution (FDR < 0.2) (Fig. 1d, Fig. 4b and Supplementary Fig. 5).

**Figure 5:**
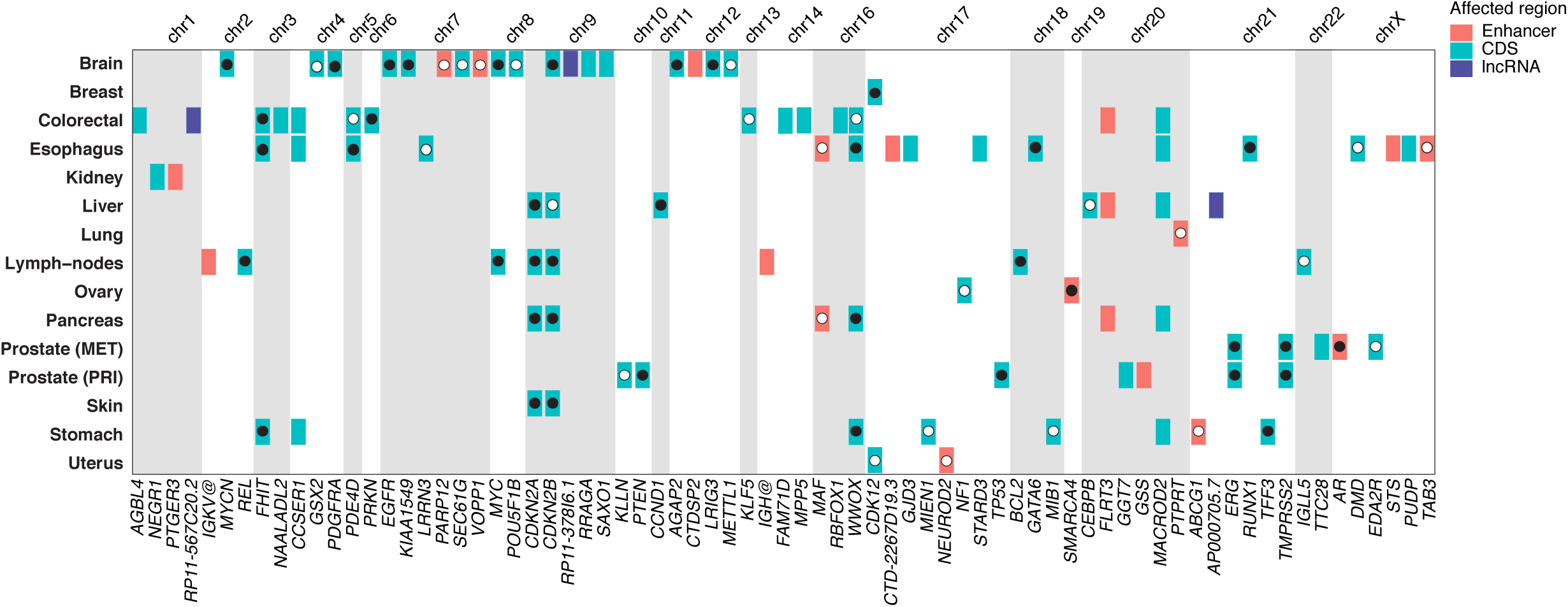
Driver candidates predicted for each cancer cohort. In green are the genes affected in the coding sequence and in red are the genes impacted at their enhancers. Genes marked with a black dot are the ones previously reported in the corresponding cancer type. Genes marked with a white dot are the ones previously reported as cancer genes but in a different cancer type than our prediction. The IGH@ locus contains 4 enhancers. The entire list of candidates is shown in Supplementary Table 5.

We identify 79 significantly recurrent peaks that potentially correspond to positively selected loci (Supplementary Table 3). The peak summits corresponding to putative driver candidates range from 179 bp to 10.71 Mb, with a median of 822.96 kb, highlighting the strength of our approach to capture breakpoint clustering over varied genomic lengths (Fig. 4c). The number of significant peaks ranges from 0 in bone cancer to a maximum of 13 in colon cancer (Supplementary Fig. 6 and Fig. 4c). Upon comparison with PCAWG results [5], we found that the novel peaks identified as significant in our analysis tend to be more cancer-type specific. Among the 42 peaks that overlap with regions of PCAWG candidates (Supplementary Table 4), 16 are cancer-type specific, while 26 were found in multiple cancer types. Conversely, 32 peaks have no overlap with PCAWG results: 25 of those peaks are cancer-type specific and 7 peaks were found in multiple cohorts. Thus, there is a significant enrichment of cancer-type specific candidate peaks identified by our approach relative to PCAWG (chi-squared test pvalue=0.0006) and further demonstrates the power of tissue-specific analysis.

**Figure 6:**
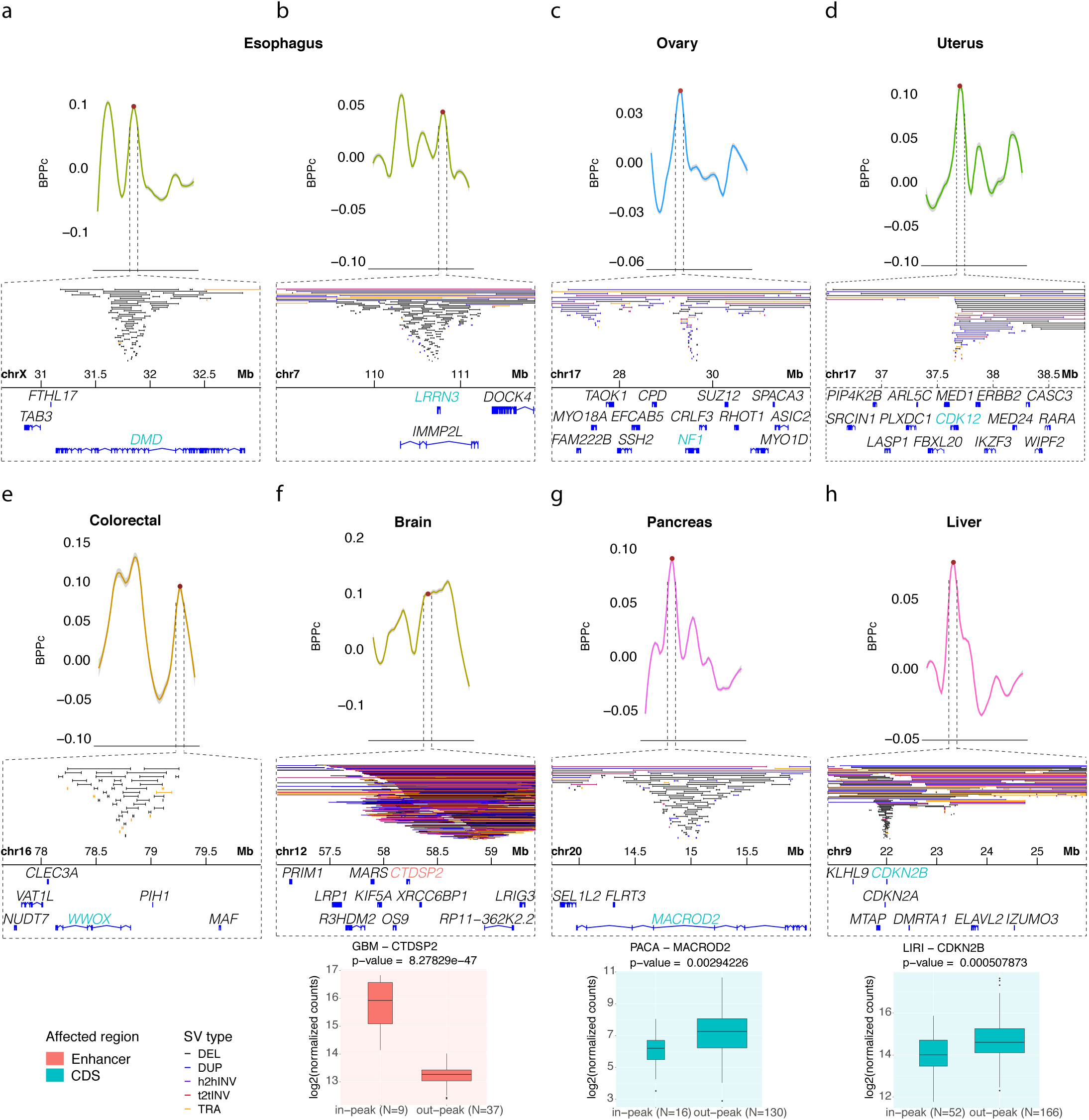
Zoomed in plots of the genome location of the significantly rearranged peaks that show the breakpoint clustering at regions of novel candidates with previous reports in other cancer types. The different types of SV are color coded as (DEL black, DUP blue, h2hINV purple, t2tINV red, TRA orange), (**a**) *DMD* in a peak on chrX in esophagus cancer, (**b**) *LRRN3* in a peak on chr7 in esophagus cancer, (**c**) *NF1* in a peak on chrl7 in ovarian cancer, (**d**) *CDK12* in a peak on chrl7 in uterine cancer, (**e**) *WWOX* in a peak on chrl6 in colorectal cancer, (**f**) *CTDSP2* in a peak on chrl2 in brain cancer, (**g**) *MACROD2* in a peak on chr20 in pancreatic cancer, (**h**) *CDKN2B* in a peak on chr9 in liver cancer. The boxplots show the differential expression between samples with and without SVs in the peak region. GBM, glioblastoma multiforme; PACA, pancreatic cancer; LIRI, liver cancer. In green are the genes affected in the coding sequence. In red are the genes impacted in their enhancer regulatory region.

Next, we predicted the functional elements that are the most likely targets of positive selection by computing the element rearrangement scores (*RS*_*E*_) for all protein-coding exons, long non-coding RNAs (lncRNAs), enhancers and CTCF-insulators in the 79 significantly recurrent peaks (Figs. 1d, 1e, Supplementary Fig. 6 and Supplementary Table 5) (Methods). The highest *RS*_*E*_ values point to elements impacted by the largest number of SVs in the maximum number of samples within the entire peak. Across all the analyzed cancer cohorts, we identified 53 coding genes, 24 enhancers of 17 other genes and 3 lncRNAs as the most likely targets (Fig. 5) (Methods). Our results are validated by known cancer genes in different cohorts. For example, *TMPRSS2*–*ERG* fusion [31, 32], and *PTEN* and *TP53* deletions [33, 34] in prostate cancer. *AR* enhancer is significantly affected only in the metastatic prostate cancer cohort, which is consistent with the development of the disease [33-35]. We also fnd *EGFR, MYCN* and *MYC* in brain cancers [36, 37]; *BCL2, MYC* and the loci of *IGH* translocations in lymph nodes cancer [38-40]; *CDK12* in breast cancer; *CCND1* in liver cancer; and *RUNX1, GATA6* and *PDE4D* in esophageal cancer (Fig. 5). Besides these known cancer genes in specific cohorts, we also identified several others with known roles in multiple cancers (Fig. 5). Interestingly, most of these common candidates are large cancer genes within fragile sites, such as *FHIT, WWOX, CCSER1, IMMP2L, CDKN2A* and *CDKN2B* [26, 41-44].

Overall, out of the 73 genes whose exons or enhancers are identified as putative drivers, 47 are known cancer genes (Fig. 5). Genes identified as potential drivers in our analysis that are known to be oncogenic in another cancer type [45, 46] are of high interest since they are often therapeutic targets under investigation (Fig. 6). For example, *DMD* is known to be a cancer gene [45] in human myogenic tumors, such as gastrointestinal stromal tumors [47], rhabdomyosarcoma [48] and leiomyosarcom [49], and we fnd it is a putative driver gene in esophageal cancer where 54% of the cohort carries SVs in this region (Fig. 6a). We also identified *LRRN3* as a driver candidate in esophageal cancer with SVs in 20.7% of the samples (Fig. 6b). Other members of the *LRRN* gene family have been found as drivers in multiple cancer types, including neuroblastoma and gastric cancer [50], and the role of *LRRN3*, in particular, has been studied in fibrosarcoma [51]. As shown in Fig. 6, clear peaks in *BPpc* correspond to tight clustering of breakpoints allowing us to confidently pinpoint these genes as putative drivers.

Another interesting result is our finding in ovarian cancer of a region affected in 25% of the cohort where our method points to *NF1* as a potential driver (Fig. 6c). Previous studies have shown the importance of *NF1* in several other cancers including glioblastoma, melanoma, breast and lung cancers [36, 52-54]. The gene *CDK12* is a candidate within a region affected in 14.9% of the uterine cancer cohort (Fig. 6d). *CDK12* is a well-studied target in other female reproductive cancers, such as ovarian and breast [55], and it has also been studied in stomach and prostate cancers [20, 56]. In colorectal cancer, we predict *WWOX* as a putative driver in a region that shows SVs in 25% of the samples (Fig. 6e). *WWOX* is within a known fragile site and has been reported in several studies of different tumor types including breast, prostate, lung, esophagus, cervical, ovarian and bladder cancers [57-61].

Limited availability of RNA-seq data prevented analysis of impact on gene expression for many candidates (Supplementary Table 6). Among the cohorts with sufficient sample sizes for RNA-seq analysis, we find that an enhancer of *CTDSP2* is a putative driver in brain cancer (glioblastoma multiforme) and the SVs are associated with its differential expression (Fig. 6f). *CTDSP2* is known to be important in other cancer types [62, 63]. The genes *MACROD2* in pancreatic cancer (Fig. 6h) and *CDKN2B* in liver cancer (Fig. 6g) also show significant differential expression between samples with SVs at these loci relative to those without SVs.

These findings highlight the importance of both the tissue-specific and clustering-based approaches used in our method to capture the significantly rearranged regions in different cancers. Notably, while breakpoint clustering clearly points to these specific candidates as seen in Fig. 6, breakpoint density over fixed bins would require different bin sizes to capture the ones with maximum density. This is because using a fixed-size window for all these instances will likely generate several bins with high density without clearly pointing to the most probable target element of selection.

### CSVDriver: Computational tool to identify SV drivers from whole-genome sequences

The computational approach developed in this work to identify SVs and functional elements under positive selection in cancer whole-genomes is implemented in CSVDriver, Cancer Structural Variation Driver, a user-friendly shiny app tool (https://github.com/khuranalab/CSVDriver). The input for the tool is SV calls and researchers can provide the tissue-specific co-variates data or use existing datasets to run the tool on their cancer cohort/s. Besides the list of functional elements that are putative drivers in a given cohort, the app provides the graphical visualization of the GAM results including the analysis of non-linearity and covariates interaction.

## Discussion

One of the major challenges in cancer genomics is the accurate estimation of the expected heterogeneous distribution of passenger variants. This distribution represents the null background, which can be used to identify loci that exhibit significantly recurrent variants likely due to positive selection. While this problem has received considerable attention for SNVs, studies for tissue-specific neutral models of the genomic distribution of SVs are lacking. We find that a GAM is able to describe the breakpoint proximity distribution of SVs in cancer genomes with the explained deviance ranging from 10% in lung to 57% in lymph-nodes cancer. The use of a GAM allows us to interpret the results graphically and provides estimates for both the magnitude and statistical significance of the contribution of each feature. We find that the 3D chromosomal conformation plays an important role in the genome-wide distribution of SVs for most cancer types with TAD-recurrence, TAD-B recurrence and their interaction terms with other covariates contributing significantly to the model.

Our method is able to identify the known cancer drivers and nominate novel candidates as those that exhibit higher breakpoint proximity than expected by random chance, such as *DMD* and *LRRN3* in esophageal cancer, *NF1* in ovarian cancer, *CDK12* in uterine cancer and *WWOX* in colorectal cancer. We note that the pan-cancer analysis by PCAWG allowed the identification of regions that are not likely to gain significance in single cancer analyses due to limited cohort sizes but failed to identify many novel regions that gain significance in our tissue-specific analysis. While our method is able to identify 24 enhancers as putative drivers, they are usually in the vicinity of other coding exons with similarly high element rearrangement scores and further functional validation will be needed to decipher their role in tumorigenesis. However, one prominent example supported by RNA-seq data is *CTDSP2* enhancer that is a candidate driver in glioblastoma.

Finally, while we could identify the features that contribute significantly to the breakpoint proximity curve, we find that the relationships are usually tissue-specific, complex and non-linear, often forbidding straightforward interpretations. Furthermore, while we focused our analysis on the peaks in *BPpc*, the valleys could potentially provide insights about the loci showing depletion of breakpoints due to negative selection in future studies.

## Methods

### Cancer somatic structural variations data

All somatic SVs were obtained from the PCAWG project Working Group 6 consensus calls [1, 18]. We selected the cohorts based on the median number of SVs per sample aiming to get data representative of the cancer types more impacted by genomic rearrangements. It covers 15 different organs systems including 32 distinct histological cancer subtypes. They are Skin: MELA (Melanoma), SKCM (Skin Cutaneous melanoma); Pancreas: PACA (Pancreatic Cancer), PAEN (Pancreatic Endocrine Neoplasms); Lymphatic system: DLBC (Lymphoid Neoplasm Diffuse Large B-cell Lymphoma), MALY (Malignant Lymphoma); Esophagus: ESAD (Esophageal Adenocarcinoma); Lung: LUAD (Lung Adenocarcinoma), LUSC (Lung Squamous cell carcinoma); Uterus: UCEC (Uterine Corpus Endometrial Carcinoma); Ovary: OV (Ovarian Cancer); Breast: BRCA (Breast Cancer); Brain (central nervous system): PBCA (Pediatric Brain Cancer), LGG (Brain Lower Grade Glioma), GBM (Brain Glioblastoma Multiforme); Bone: SARC (Sarcoma), BOCA (Bone Cancer); Colorectal: COAD (Colon Adenocarcinoma), READ (Rectum Adenocarcinoma); Kidney: KICH (Kidney Chromophobe), KIRC (Kidney Renal Clear Cell Carcinoma), KIRP (Kidney Renal Papillary Cell Carcinoma), RECA (Renal clear cell carcinoma); Liver: LICA (Liver Cancer), LIRI (Liver Cancer - RIKEN), LIHC (Liver Hepatocellular carcinoma), LINC (Liver Cancer - NCC); Stomach: GACA (Gastric Cancer), STAD (Gastric Adenocarcinoma); Prostate: EOPC (Early Onset Prostate Cancer), PRAD (Prostate Adenocarcinoma). Particularly for prostate cancer, we additionally collected SVs from a cohort of 124 metastatic samples (37,9605 SVs) obtained from calls reported in [19] and [20]. These datasets provide information from 2,382 high-quality WGS cancer donors allowing us to analyze 324,838 high confidence unique somatic SVs. We analyzed SVs in autosomes and chromosome X only. The genome reference used is hg19 build. The details are shown in Supplementary Table 1.

### Genomic feature annotations and tissue-specific epigenomic covariates

We use several genome features as the *BP* covariates in the modeling of the expected background of *BPpc*. They include tissue-specific chromatin state marks from Roadmap Epigenomics Mapping Consortium [23], which annotates genomic regions based on histone modifications and chromatin DNA accessibility [64]. We used the average RT signal per 1Mb genomic bins across eight different cell types (liver, breast, brain, lymphoma system, skin, blood, lung and prostate) as described in the CNC-Driver method [11]. The datasets were collected from ENCODE and constitute wavelet-smoothed Repli-seq data [25]. The GC content was computed in windows of 101 bp centered in each BP using the function ‘GCcontent’ from the R Package ‘biovizBase’ [65]. The gene density was computed per 1 Mb window using the function kpPlotDensity from the R Package kpPlotDensity [66]. The chromosomal Fragile Sites (FS) annotation was obtained from the database HumCFS [26].

We included two higher-order structural 3D chromosomal conformations, TADs and LADs, in the set of covariates. For TADs, we use the recurrence of the regions as well as their boundaries (TAD-B) annotated across 37 datasets obtained from the 3D Genome Browser [24]. We also classify each sub-segment of TAD that shows different recurrence by computing the coverage of all 15 chromatin marks within every TAD segment, similar to the approach used in [16] (Supplementary Fig. 7). We found that three principal components explain most of the variance of this coverage (Supplementary Fig. 8). Thus, we grouped the TAD segments in three clusters according to the coverage of chromatin marks. The three clusters are quiescent/heterochromatin, low-active and active. We confirm that the clustering is robust across the 16 cohorts analyzed in this study (Supplementary Fig. 9). More details of the sources of genomic feature annotations are shown in Supplementary Table 2.

The putative functional effect for each significantly rearranged locus is predicted by annotating the potential drivers on the basis of the impact of the SVs breakpoints on the coding and noncoding elements within these regions. For the coding drivers, we use the subset of protein-coding genes extracted from the comprehensive gene list obtained from the Genecode Release 29 (GRCh37). For the non-coding elements, we use the active tissue-specific cCRE marks gathered from ENCODE 3 [25].

### CSVDriver workflow

CSVDriver aims to identify cancer-driving rearrangement events by analyzing the focal trend of breakpoint clustering.

#### Input and preprocessing of data

The method takes as input the data of cancer somatic SVs calls and analyzes the combined impact of all rearrangement types including insertions (Ins), deletions (Del), inversions (Inv) and translocations (Tra). The expected file format is SV-table containing 12 columns named as: (cohort_code, donor_id, variant_type, sv_id, chr_from, chr_from_bkpt, chr_from_bkpt, chr_from_strand, chr_to, chr_to_bkpt, chr_to_bkpt, chr_to_strand). Each row defines a single SV by the cohort (cancer type), the donor ID, the SV type (Ins, Del, Inv, Tra), the SVs unique ID and the genome coordinate [chr, start, end, strand] for both rearrangement breakpoints (from, to). CSVDriver checks SVs from all donors and decomposes this SV-table into a BP-table sorted by genomic position per chromosome.

#### Establishing the observed breakpoint proximity-curve (*BPpc*)

The method bases the SV analysis on the description of the genome-wide distribution of breakpoint proximity taking all unique breakpoint coordinates for each distinct sample in the given cohort. Thus, if a breakpoint coordinate is present in two samples, it is represented twice. Then we sort all breakpoints based on their coordinates and obtained an ordered list of breakpoints encompassing all samples, each one represented by *BP*_*i*_, where *i* is the ordered index. Next, we annotate each *BP*_*i*_ by computing their neighbor reachability (*BPnr*_*i*_,) defined as the average genome distance to reach the two adjacent breakpoints:

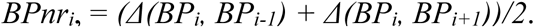

Thereupon we define the breakpoint proximity (*BPp*_*i*_) as the normalized reverse scale, applying logarithmic transformation: *BPp*_*i*_ *= -log10(BPR*_*i*_*+1)*. Then, we compute the breakpoint proximity curve (*BPpc*), which is a smooth curve resulting from the nonparametric local polynomial regression (LOESS) fitted to the *BPnr*_*i*_ values (Fig. 1A). This curve shows the trend of focal clustering of the breakpoints because the fitting result is weighted toward the nearest surrounding values. The span argument (α = 0.2) controls the size of the near surrounding. It reflects the interval as a proportion of the total breakpoints and regulates the grade of smoothness in the resulting *BPpc*. This curve depicts the breakpoint genomic distribution and captures the regions of high and low proximity between breakpoints. Chromosomes with a low load of breakpoints that show no clear trend of clustering will not reflect a reliable signal. Therefore, we do not consider them. That is the case for small chromosomes (e.g. chr21, chr22) in some cancer cohorts.

#### Modeling the expected BPpc by using generalized additive model (GAM) with tissue-specific genomic covariates

The model is conceived to expand the linear regression analysis of genomic covariates by introducing the capacity to investigate the potential non-linear relationships between genomic covariates and the distribution of breakpoints. Additionally, it accounts for the contribution of the covariates’ interaction to the model. Thus, CSVDriver models the expected background *BPpc* using GAM, a parametric regression method, which models the *BPpc* (dependent variable) with respect to tissue-specific data of genomic covariates (predictors or independent covariates).

GAM is a flexible extension of generalized linear models (GLM). Using a GAM, we can fit a linear model, which allows us to consider either linear or non-linear contributions of the genomic covariates to the model of *BPpc*. To fit the GAM, CSVDriver uses the R package ‘mgcv’ version 1.8-28 [67, 68]. The values of the observed *BPpc* for each cohort fit gamma distribution (Supplementary Fig. 10). Therefore, the model assumes each expected *BPpc* to be generated from a gamma distribution for the identity link function of the response. The model follows the equation:

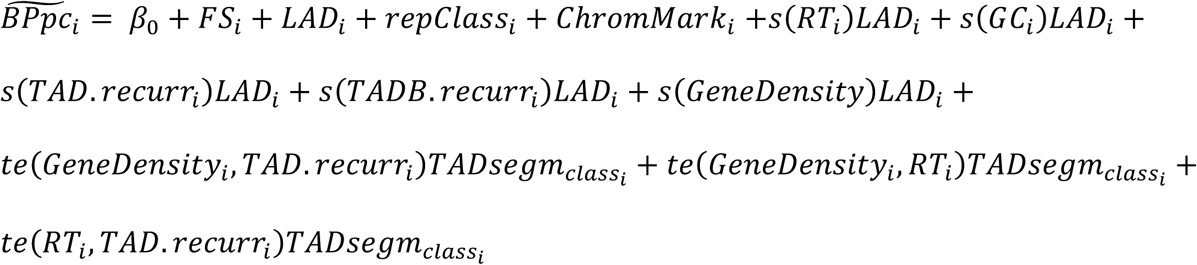

where i=1,…,N (total number of breakpoints), 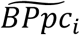 is the expected breakpoints proximity, *β*_0_is the intercept and the remaining terms are the genomic covariates used as predictors. The covariates FS, LAD, repClass, and ChromMark are modeled as linear factorial terms. The thin plate regression spline smooth function (s) can describe the non-linearity in the contribution of the covariates (RT, GC, GeneDensity, TAD, and TAD-B recurrence) and for each one the model gets the interaction with the status of the factor LAD. The model investigates the main effects of the predictors as well as the effects of the tensor product interaction (te) for the interplay between GeneDensity and TAD recurrence; GeneDensity and RT; and TAD and RT. This accounts for the status of TAD-segment class.

Nonetheless, a high number of interaction terms increases the chance of over-fitting. Furthermore, GAM can be computationally expensive to reach convergence. Consequently, we try to balance the complexity of the model and its ability to explain the deviance by covariate interactions reasonably by including the interaction only of covariates that contribute at least 10% in at least one cohort in single predictor models.

#### Computing the adjusted *BPpc*

The goal of CSVDriver is to capture the loci where breakpoint clusters potentially arise due to selective pressure unlike the clusters associated with neutral non-selective forces. Therefore, the adjusted curve 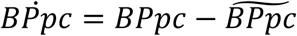 represents a corrected BPpc where regions with values close to the 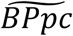 will be considered expected biases. The signal in the y-axis depicts how close the breakpoints are in a given region (Fig. 1c). The positive values represent regions where the breakpoints are closer than expected while regions with negative values are loci where the breakpoints are sparser than expected from the GAM. The values around zero in the adjusted curve 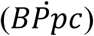 correspond to regions of observed BPpc close to the expected one.

#### Detecting the significantly rearranged peaks in the *BPpc*

The method takes the regions corresponding to the top 25% of the summit of each peak, which represent the regions of local maximum breakpoint clustering. Then we identify the loci that potentially correspond to the positively selected regions (Fig. 1d). Each peak is described by their peak recurrence score (*PRs*) defined as:

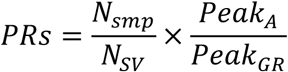

where *Nsmp* is the number of unique samples in the peak, *Nsv* is the number of unique SVs, *Peak*_*A*_ is the area under the peak and *Peak*_*GR*_ is the genome range of the peak. Thus, *PRs* is the highest for peaks where many samples (high *Nsmp*) contribute to create tight clusters of breakpoints (high *Peak*_*A*_) over narrow regions (low *Peak*_*GR*_). The score is square root transformed to reduce the dispersion in the values while keeping the trend of interest. We take into account the inherent limitation of the background model and the differential explained deviance per tissue type (Fig. 2a). Consequently, the approach does not use any absolute pre-established threshold, instead, it considers the cohort-specific distributions of all *PRs* (Fig. 4b density distribution) as the expectation of the combined effect of several processes. Next, for each cohort, we identify peaks with *PRs* significantly higher (Fig. 4b, QQ-plots) than the fitted gamma distribution theoretical density using the ‘fitdistrplus’ R Package [69]. After controlling the false discovery rate (FDR) [70] for multiple hypothesis testing, the significant loci were defined as those with FDR < 0.2.

#### Detecting the driver candidates within the significantly rearranged peaks

The potential functional effect of the significantly rearranged regions for each cancer is directly associated with the effects on coding and non-coding elements by their deletion, disruption or relocation. CSV-Driver determines the functional elements that are the most likely targets of positive selection within each significantly rearranged peak.

For each functional element (protein-coding exons, lncRNA, enhancer, CTCF-insulator) located at a significantly rearranged peak, the method computes the element rearrangement-score (*RS*_*E*_)

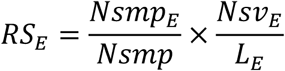

where *Nsmp*_*E*_ is the number of unique samples that have SVs overlapping the element, *Nsmp* is the number of unique samples in the peak, *Nsv*_*E*_ is the number of SVs overlapping the element and *L*_*E*_ is the length of the element. High *Nsv*_*E*_ values could be due to multiple alleles affected or multiple clonal/sub-clonal cell populations with SVs at the given locus. Normalization by *L*_*E*_ takes into account that longer elements are more likely to have more SVs randomly. *RS*_*E*_ is a metric of the relative selective pressure acting upon elements within the significantly rearranged region. It takes into account the relative recurrence in samples and the number of SVs that impact the element normalized by the length of the element. Due to the larger genomic length of genes and different mechanisms of SVs that can impact their function, i.e. (a) amplification, deletion, relocation or (b) gene fusion, CSVDriver computes two rearrangement scores *RS*_*E*_ for coding genes under significant peaks. One score accounts for the number of SVs that engulf the entire gene and the other accounts for the number of breakpoints that fall within the gene’s body. The genes with the highest *RS*_*E*_ are the most likely targets of positive selection.

The method reports the element rearrangement scores for the highest scoring elements of all types (protein-coding exons, enhancer or lncRNA) at a given peak (Supplementary Fig. 6 and Supplementary Table 5). While at a specific locus altered by SVs, it is fair to assume that affected coding exons of genes are more likely to have greater impact than the non-coding elements, CSV-Driver allows the analysis of the potential importance of all possible driver elements. One peak may contain multiple driver elements which can represent alternative paths of disrupted regulation, or even some subgroup of samples with slightly different, yet related genomic drivers. In the current analysis, if a peak contains multiple elements of different type, all elements with highest rearrangement scores are reported in Supplementary Fig. 6 and Supplementary Table 5. However, if one of those elements is or is associated with a known cancer gene, we consider it as the most likely candidate in a given peak shown in Fig. 5.

#### Results report

For each input cohort, CSVDriver reports the graphics of the *BPpc* annotated with the driver candidates (coding genes and non-coding elements) at each significantly rearranged peak (Supplementary Fig. 6). It also provides the summary tables with the catalog of drivers (Supplementary Table 5). The tool also provides the set of plots resulting from GAM covariates modeling to analyze the non-linearity of relationships between the genomic features and the genomic distribution of breakpoints (Supplementary Fig. 3)

### Verification of the driver candidates

We verified our results of driver candidates using CancerMine [45] and COSMIC [46]. In addition, for the significantly rearranged peaks that have enough samples with RNA-seq, we check for the significant differential expression between samples with SVs at the given locus relative to the samples without SVs.

## Acknowledgement

This work is supported by NIH grant R01CA218668 to E.K.

## Author Contributions

E.K. and A.MF. conceived and designed the project. A.MF. implemented the method and performed the data analysis. A.D. performed the RNA-seq differential expression analysis. A.MF. and E.K. wrote the manuscript. E.K. supervised the project.

